# Ultrasensitivity of Microtubule severing rate on the concentration of free Tubulin

**DOI:** 10.1101/2022.04.26.488783

**Authors:** Chloe E. Shiff, Jane Kondev, Lishibanya Mohapatra

## Abstract

Cytoskeletal structures aid in cell polarization, motility, and intracellular transport. Their functions are predicated on the rapid turnover of cytoskeleton proteins, which is achieved by the coordinated effort of multiple regulatory proteins whose dynamics are not well understood. In-vitro experiments have shown that free tubulin can repair nanoscale damages of microtubules created by severing proteins. Based on this observation, we propose a model for microtubule severing as a competition between the processes of damage spreading and tubulin-induced repair. Using theory and simulations, we demonstrate that this model is in quantitative agreement with in vitro experiments. We predict the existence of a critical tubulin concentration above which severing becomes rare but fast, and hypersensitive to the concentration of free tubulin. Further we show that this hypersensitivity leads to a dramatic increase in the dynamic range of steady-state microtubule lengths, when lengths are controlled by severing. Our work demonstrates how synergy between tubulin and severing proteins can lead to novel dynamical properties of microtubules.

## Introduction

Cells contain a cytoskeleton that aids in various cellular processes such as maintaining cell polarization, motility, and transport. The cytoskeleton is highly dynamic – it is composed of various structures that are constantly built and dismantled to assist in various functions. For example, the single-cell alga *Chlamydomonas* uses microtubule-based flagella for motility (Marshall et al., 2005; Song and Dentler, 2001). Microtubule-based fibers in the mitotic spindle shorten for chromosome separation during mitosis and meiosis (Bennabi et al., 2016; Dumont and Mitchison, 2009; Hyman and Karsenti, 1996).

Such dynamic changes in cytoskeletal structures require a rapid turnover which involves an exchange of their molecular building blocks, with the same molecules that are freely diffusing in the cytoplasm. Remarkably, the time scales associated with the dynamics of proteins filaments in vivo are quite different from those measured in vitro. While actin networks within cells are disassembled on the order of seconds, the process takes a few minutes in vitro (Kuhn and Pollard, 2005; Miyoshi and Watanabe, 2013; Pollard, 1986; Watanabe and Mitchison, 2002). Similarly, the polymerization rate of tubulin in vivo is about five to tenfold higher than in vitro at a similar concentration of free tubulin (Desai and Mitchison, 1997). Further, microtubules are known to switch between phases of slow growth and rapid shrinking by a process known as dynamic instability, which has been studied extensively in vitro (Desai and Mitchison, 1997). The rates of switching from shrinking to growth and vice versa (called rates of rescue and catastrophe, respectively) are sensitive to, and are believed to be modulated by several microtubule-associated proteins in vivo (Desai et al., 1999; Gardner et al., 2013; Geisterfer et al., 2020; Goshima et al., 2005; Tournebize et al., 2000). Turnover rates can also be different at different stages of the cell cycle – for example, microtubule turnover rates measured in Xenopus extracts range from 3 min to 20s during interphase and mitosis, respectively(Hyman and Karsenti, 1996). While much is known about the biochemical processes that facilitate the turnover of these structures in isolation, it is still not clear how different cytoskeleton-associated proteins work together to promote filament turnover in cells.

Recently, several studies have addressed this question by combining multiple factors with cytoskeletal filaments in vitro and have reported novel synergistic results that are not seen when studying each component individually. Shekhar et al. report that the cofilin-induced depolymerization rate of actin filaments increases 200-fold in the presence of cyclase associated protein/Srv-2(Shekhar et al., 2019). Similar studies have been done using microtubules with a recent study finding that collective effects of microtubule-associated proteins like MCAK and XMAP215 can lead to microtubule treadmilling(Arpağ et al., 2020). Inspired by yet another set of recent experiments, we consider the interplay between the free pool of tubulin dimers and microtubule severing proteins, which fragment protein filaments(Vemu et al., 2018).

Free tubulin is known to affect microtubule dynamics by modulating rates of nucleation and polymerization (Geisterfer et al., 2020; King and Petry, 2020; Ohi et al., 2021; Woodruff et al., 2017). Notably, using FRET, it was shown that a local release of tubulin promoted microtubule extensions into lamellipodia, demonstrating a feedback between filament bound and free tubulin pools(Van Geel et al., 2020). However, there are also reports of free tubulin affecting microtubule disassembly via its interactions with proteins such as kinesin-13 (Desai et al., 1999; Walczak et al., 2013), kinesin-8 (Arellano-Santoyo et al., 2017; Varga et al., 2009) and severing proteins (Kuo et al., 2019b; Vemu et al., 2018). Here we theoretically describe the effect of free tubulin on microtubule severing via a recently discovered process of repair of damages to the microtubule lattice by the severing proteins(Vemu et al., 2018).

Severing proteins can play a key role in the reorganization of the cytoskeleton by breaking down existing structures and making more monomers available for use in new structures (Kuo and Howard, 2021). Recent studies have found that severing proteins can seed new filament growth by releasing filaments from nucleation sites and allowing their transport within the cell (Sharp and Ross, 2012). Katanin and spastin are well characterized examples of severing proteins for microtubule-based structures, while cofilin and Srv-2 sever actin-based structures (Balcer et al., 2003, p. 2; Chaudhry et al., 2013; Díaz-Valencia et al., 2011; Johnston et al., 2015, p. 2; Kuo et al., 2019b). Katanin is thought to play two functions in mitosis and meiosis – to amplify the number of microtubules in the spindle and to uncap microtubule plus ends from the kinetochore and enable depolymerizing kinesins to target microtubules (Sharp and Ross, 2012). Inhibition of katanin results in abnormally long mitotic spindle length in *Xenopus* extracts (Loughlin et al., 2011). Severing proteins play an important role in neurons as well (Karabay et al., 2004). It has been proposed that katanin stimulates axon growth by releasing microtubules from centrosomes and severing them into shorter segments that can be transported along longer microtubules down the axon to seed new microtubule growth (Ahmad et al., 1999). Mutations in *Drosophila* spastin reduce dendritic arborization (branching) (Jinushi-Nakao et al., 2007), suggesting that spastin seeds microtubule growth within newly forming regions of the dendrite. Additionally, by using in vitro systems and theory, many studies have also analyzed the effect of severing proteins on the length of cytoskeletal filaments (Elam et al., 2013; Kuo et al., 2019b, 2019a; Mohapatra et al., 2016; Roland et al., 2008; Tindemans and Mulder, 2010).

Thus far, most studies have focused on the effect of severing proteins in isolation, but recently experiments have examined the interplay of these proteins in presence of free tubulin. Electron microscopy images of microtubules have revealed the mechanisms by which severing proteins induce a break in the microtubule. One study reported that once severing proteins land on a microtubule, they create nanoscale damages(Vemu et al., 2018). This study found that not all of the damage sites proceeded to a severing event, as free tubulin could repair the damage in some cases. Furthermore, a few in-vitro studies have reported that severing proteins enhance the rate of rescue in microtubules undergoing dynamic instability (Kuo et al., 2019b, 2019a; Kuo and Howard, 2021; Vemu et al., 2018). At this point, it is unknown whether severing proteins have the same effect in vivo, where they may interact with other microtubule-associated proteins.

Inspired by these observations (Vemu et al., 2018), we propose a model for microtubule severing that incorporates the newly discovered repair process by free tubulin. Using theory and simulations, we show that this new model quantitatively agrees with in vitro experiments and makes predictions for new in vitro experiments. Significantly, we predict the existence of a critical tubulin concentration above which severing becomes rare but fast. This model also has several implications for the dynamics of microtubules in vivo – we show that length control via severing is far more sensitive to changes in tubulin concentrations, resulting in a dramatically expanded dynamic range of filament lengths. We also report the probability distribution of lengths and the mean length and its coefficient of variation as a function of the model parameters, which can be used to test the model. In summary, our work describes how the concerted action of tubulin and severing proteins produces novel dynamical properties of microtubules that are not seen when these proteins are studied independently.

## Results

### 1. Model of microtubule severing with repair

Inspired by recent experiments(Vemu et al., 2018), we consider that a microtubule severing event begins when a severing protein lands on a filament, binds to the side, and creates a damage spot (Fig. 1A). The fate of the damage created by a severing protein is determined by a competition between the removal and addition of tubulin at the damage site, eventually leading to severing or repair. We assume that monomers are added to the damage site at a rate proportional to the concentration of free tubulin *k*_*T*_[*T*] (*s*^−1^) and are removed from the damage site at a rate *k*_*r*_ (*s*^−1^), with each step either decreasing or increasing the size of damage. If the size of the damage reaches a size *N*_*D*_ removed tubulins, the filament is severed. On the other hand, if the size reaches zero, then the damage is repaired. In this way, the size of the damage *x* measured in monomers (i.e., tubulin dimers) is modeled as a one-dimensional random walk with absorbing boundaries at *x* = 0 and *x* = *N*_*D*_. (Fig. 1B)

**Figure 1.**
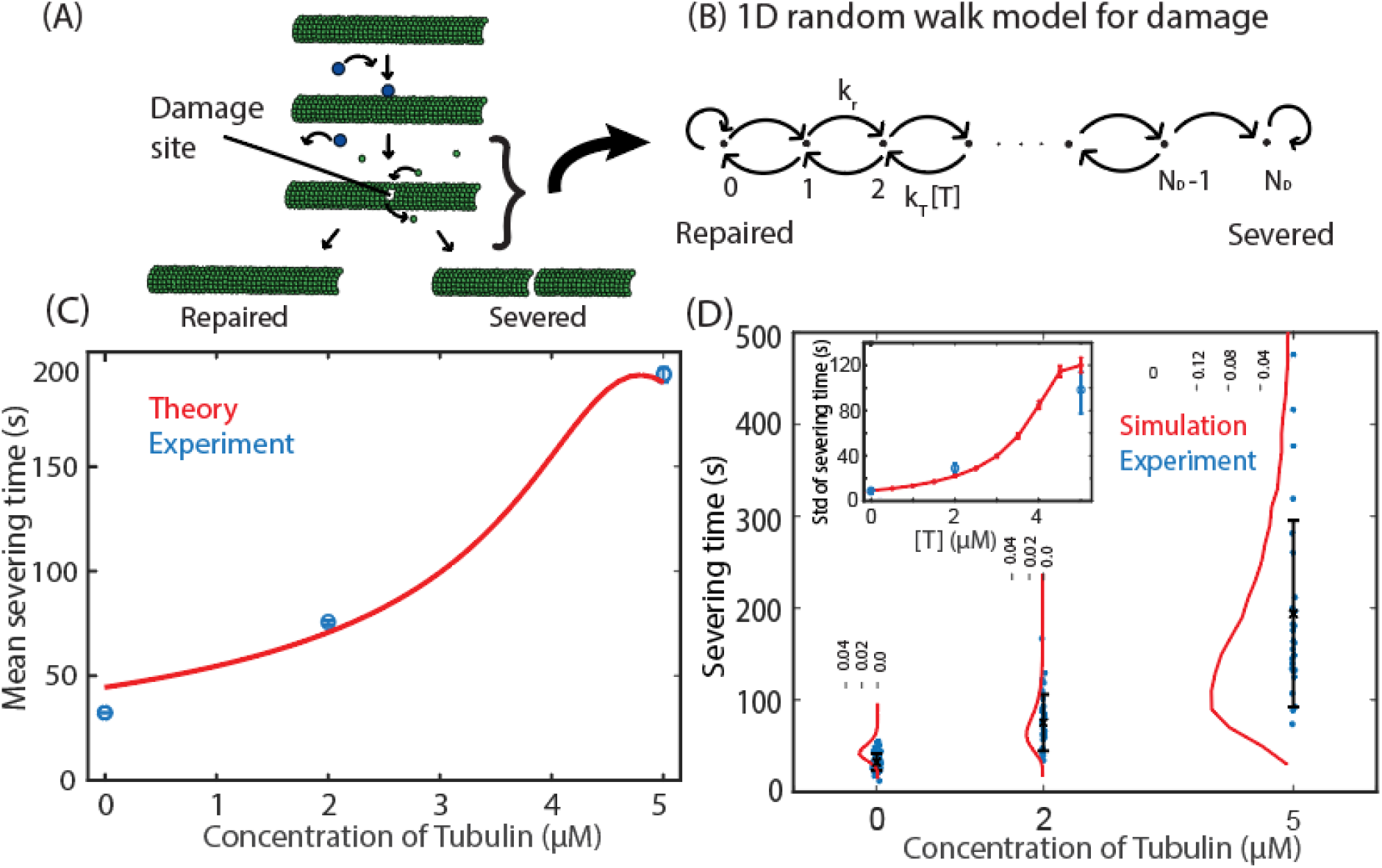
Microtubule severing with repair. (A) Severing proteins (in blue) create small damages on microtubule lattice which either get repaired by free tubulin or lose enough monomers to sever the microtubule. (B) One-dimensional random walk model for studying the competition between removal of tubulin monomers at a rate *k*_*r*_ and the addition of monomers at a rate proportional to the free tubulin concentration, *k*_*T*_[*T*].(C) We use published data (Vemu et al., 2018) for the severing times as a function of tubulin concentration to estimate parameters of the severing model, which\ are listed in Table 1. (D) The same parameters are used to compute the standard deviation (inset) and probability distribution of severing time using stochastic simulations. Each distribution was computed from 400,000 trajectories. Error bars on experimental data obtained by bootstrapping method (Section 4 in Materials and Methods). The error bars on the standard deviation from simulations show SEM calculated from 750 values, with each value computed from 20,000 trajectories. Comparison with published experimental data (Fig. A2, from (Vemu et al., 2018), in blue) is shown.

In this model of severing with repair, a damage site can either be repaired or progress to a severing event. Using our one-dimensional random walk model, we find an analytical expression for the probability of severing (described in section 2 and SI), and the mean time it would take to sever a filament as a function of model parameters (*k*_*T*_, *k*_*r*_, and *N*_*D*_). Assuming an initial damage size of one monomer, and defining the ratio of the tubulin addition and removal rate as a dimensionless tubulin concentration 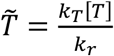, we predict that the time to sever a filament has the following form 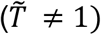

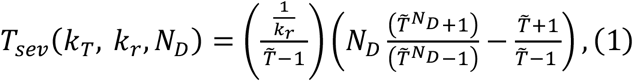

**Table 1.**
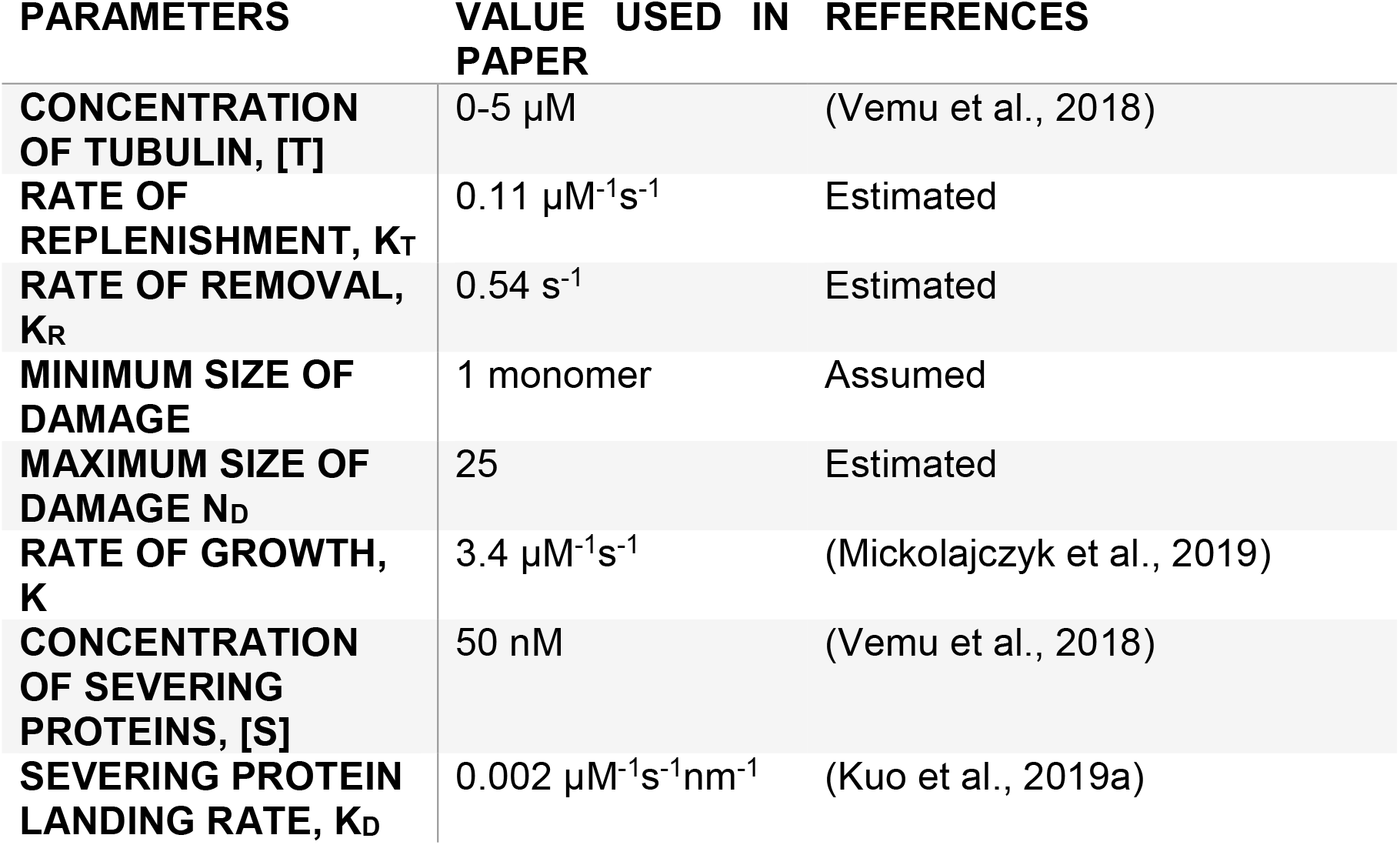
Parameters used in the models for the model of severing and length control with tubulin-induced repair. Estimated values were attained by fitting the theoretical expression for the mean severing time to experimental data in ref (Vemu et al., 2018); see Figure 1C.

In the case, 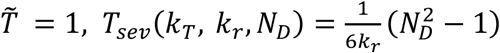

Considering *N*_*D*_ ≫ 1, equation 1 has two limits. When the rate of tubulin removal is more than tubulin addition, i.e., 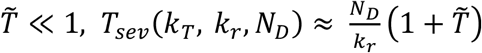 the severing time is linear in tubulin concentration. When 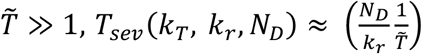, the severing time decreases as tubulin concentration increases. These limits indicate that there is a critical concentration of tubulin, 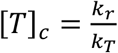, which delineates different functional dependencies of the severing time on the concentration of free tubulin. We also note that the time scale of the severing time is given by a parameter 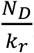, which is the mean time for completing *N*_*D*_ tubulin removal steps, each lasting 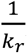, on average. Equation 1 was checked against stochastic simulation of severing with repair (described in Section 5(B) of Materials and Methods).

Next, we use published data to estimate a set of parameters for the model of severing with repair. We consider the measured probability to sever and severing time as a function of tubulin concentration published in Fig. 3D and 3H in (Vemu et al., 2018), and fit our model parameters (*k*_*T*_, *k*_*r*_, and *N*_*D*_) using method of least squares, described in Section 4 of Materials and Methods; see Figure 1C. The sen tivity of our results on the chosen parameters is shown in Fig. A1. Using the estimated parameters, listed in Table 1, we conduct stochastic simulations of the model (see section 5(A) in Materials and Methods and Figure A1 in the Appendix) to compute the distribution of severing times as a function of tubulin concentration and compare it to published data. Remarkably, just as in the data (Vemu et al., 2018), we find that the distribution of severing times is asymmetric around the maximum and skewed towards large times; see Figure 1D. A quantitative comparison between the measured standard deviation of the severing time and the predictions from our model is shown in the inset of Figure 1D.

The published experiment (Vemu et al., 2018) suggests that severing time is monotonic in the tubulin concentration up to 5 *μM*. Interestingly, our model (equation 1) predicts a non-monotonic relationship between severing time and the concentration of tubulin. We find, as reported in the study, that severing time increases as the concentration of tubulin is increased but only until a certain critical concentration of tubulin. Above this concentration, as shown in equation 1, we predict that the severing time should decrease (See Fig. 2A). As we describe next, at high concentrations of tubulin, we expect that the severing events are rare (but rapid) and almost all damages are likely to quickly be repaired by the free tubulin.

**Figure 2.**
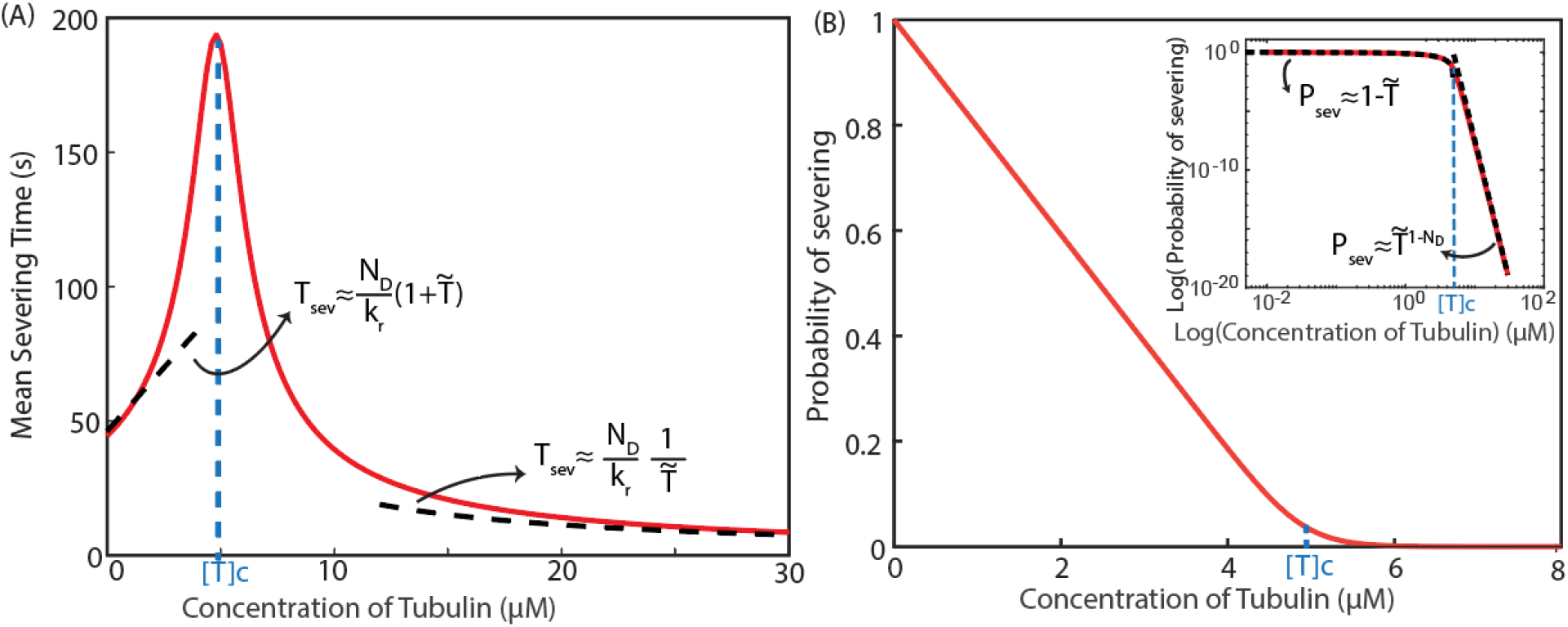
Theoretical prediction for model for microtubule severing with repair. (A)The model predicts a non-monotonic relationship between severing time and free tubulin concentration and the existence of a critical concentration of tubulin, [*T*]_*c*_, beyond which severing time decreases. (B) Probability of severing decreases as the tubulin concentration is increased (Inset: plot on a log scale). The dashed lines in both plots show the asymptotic behavior at large [*T*] ≫ [*T*]_*c*_ and small [*T*]≪[*T*]_*c*_ free tubulin concentrations. Parameters used to plot these graphs are listed in Table 1.

Note that the non-monotonicity in the severing time is a prediction of our model that can be used to experimentally test the proposed mechanism of severing with repair.

### 2. Theoretical predictions of severing with repair

Our model makes several other predictions. The probability of severing is the probability that the one-dimensional random walker reaches the *x* = *N*_*D*_ boundary before the *x* = 0 boundary:

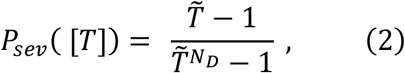

where we have chosen the initial damage size of 1, while *N*_*D*_ is the damage size that produces a severing event, and we expect *N*_*D*_ ≫ 1 (see Table 1). Again, this expression has two limits (shown in dotted black lines in Fig. 2): when 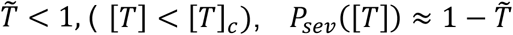, whereas when 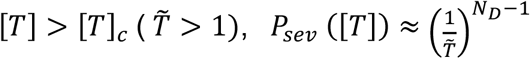. Thus, above the critical concentration, the probability of severing is very small, and damage events that lead to a severing event, are very unlikely. We confirmed our analytical expression for the mean severing time and the probability of severing as a function of tubulin concentration by stochastic simulations (See Fig. A2).

### 3. Length control in the presence of repair

The probability that a severing protein which is diffusing in solution lands on a filament is proportional to its length (Berg, 1993). Therefore, the process of severing is a length-dependent disassembly mechanism wherein longer filaments are more likely to be severed. This results in negative feedback of filament length on a growing filament and leads to control of length of cytoskeletal filaments. A number of theoretical and experimental studies (Kuo et al., 2019b, 2019a; Loughlin et al., 2011; Mohapatra et al., 2016; Roland et al., 2008; Tindemans and Mulder, 2010) have explored this aspect of severing. Given the recent observation of tubulin-induced repair (Vemu et al., 2018), we next study the implications of this repair process on microtubule length control by severing. We consider a situation where a microtubule filament undergoes growth with a rate proportional to the free tubulin concentration and severing, where severing competes with tubulin-induced repair, as described above. We make a simplifying assumption, namely we ignore the time it takes for the severing event to complete once the initial damage is created. We also assume that microtubules are stabilized and are not undergoing dynamic instability. Later in the Discussion and in the Appendix, we examine these assumptions and show that they do not qualitatively change our results about the mean steady-state microtubule lengths (see Sections A1 and A2 in Appendix, and Figs. A3 and A4), while allowing us to obtain analytic results for the steady state microtubule length. We compare our results of the severing with repair model to the severing model without repair, as considered previously in (Kuo et al., 2019a; Mohapatra et al., 2016). Results of this comparison are shown in Fig. 3.

**Figure 3.**
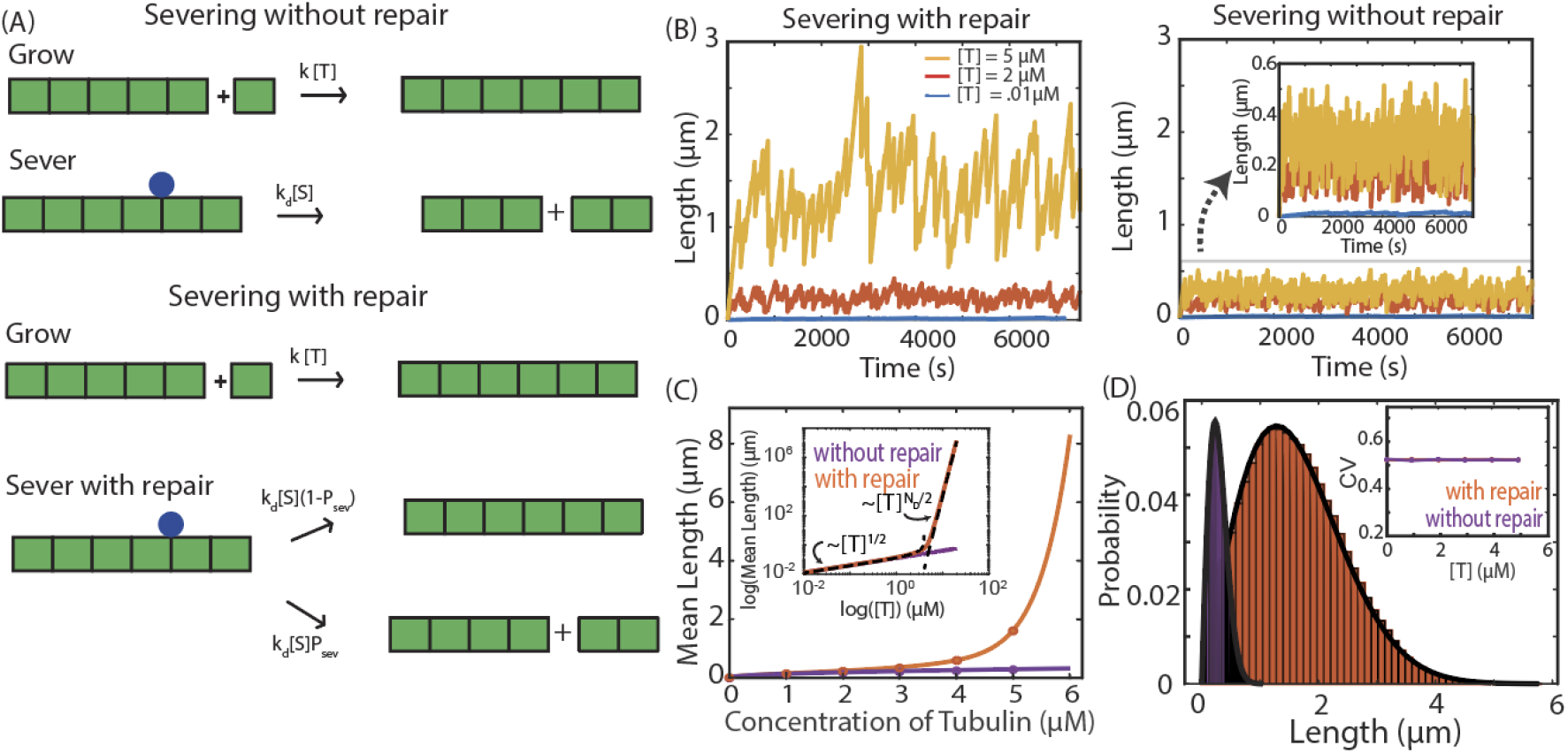
Effect of repair on filament length control via severing. (A) Schematic of the two models considered. Severing without repair: A filament grows with a rate *k*[*T*] and is severed with a rate *k*_*d*_[*S*]. Severing with repair: Filaments grow with a rate *k*[*T*] and either get severed with a rate *k*_*d*_[*S*]*P*_*sev*_ (per monomer of the filament) or repaired with a rate *k*_*d*_[*S*](1 − *P*_*sev*_) where *P*_*sev*_ (probability of severing) is calculated using equation 2. If severing occurs, the end portion of the filament (right part) is removed. (B) Dynamics of filament length (average of 3 length trajectories shown for clarity) for the two models obtained using stochastic simulations. Dynamic range of the steady state filament lengths as [T] is tuned is larger with repair than without. (C) Mean steady state length as a function of tubulin concentration (Log-Log plot inset). Dots are from simulations; lines are from analytic calculations for the model with repair (orange) and without (purple). The dashed lines represent the show the asymptotic behavior at large [*T*] ≫ [*T*]_*c*_ and small [*T*]≪[*T*]_*c*_ (D) Distribution of steady state lengths and coefficient of variation (inset) as a function of tubulin concentration, each computed from 10^6^ trajectories. Parameters in the steady-state simulations are listed in Table 1.

Using the computational scheme described in the Methods (See section 5(B)), we conduct stochastic simulations of the filament length as a function of time and analyze the process with and without the repair of severing protein-induced damage (Fig. 3A). In our simulations, individual filaments grow at a rate proportional to the tubulin concentration, *k* [T], where *k* is the second order rate constant for the tubulin binding to the end of microtubules and has the units of (*μM. s*)^−1^. Severing proteins land on the side of the filament with a rate proportional to the concentration of severing proteins [S] and to the length of the filament, *l*. For the simulations without the repair process, severing at any position on the filament is *k*_*d*_[*S*]. In the simulation with the repair process, we take *k*_*d*_[*S*]*l* to represent the rate at which damage sites are created along the length of the microtubule. Assuming this is the rate limiting process in severing, the rate of severing can be written as *k*_*d*_[*S*]*lP*_*sev*_ where *P*_*sev*_ is the probability to sever (equation 2).

For the two cases (severing without and severing with repair), we generate trajectories of length over time. In both cases, we notice that, after a steep growth phase, the filament length fluctuates around a steady-state length, but the values of range of lengths obtained for each case are different. We generate trajectories at three different tubulin concentrations and find that the mean lengths achieved by the filaments (defined as the dynamic range of filament lengths) are larger when the process of repair is included (Fig. 3B).

We also analyzed this process using theory (described in Section 6 in Materials and Methods) and calculated the mean filament length at steady state as 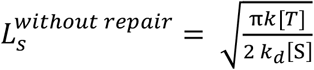 and 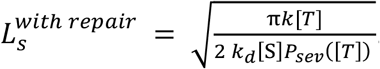. These expressions were also confirmed using simulations. As shown in Fig. 3C, while the steady state length 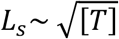 without repair, we observe two regimes with tubulin-induced repair present– when 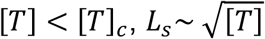 and when 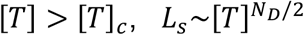, suggesting a very high sensitivity to tubulin concentration, above the critical concentration (See Section 6 in Materials and Methods). Note that our results for severing without repair 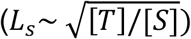 are similar to the predictions for the “no catastrophe model” in a recently published paper which studied the effect of the severing protein spastin on microtubule length (Kuo et al., 2019a). In contrast, we find that including the possibility of repair of severing events in our model predicts a new regime of microtubule growth that was not described previously.

We derive analytically (described in Section 6 in Materials and Methods), the probability distribution of the steady state lengths and compare our simulations with the theory in Fig. 3D. We also calculate the coefficient of variation (CV = standard deviation/mean, commonly referred to as the noise) of the steady-state lengths, with and without repair. Significantly, we calculate that CV doesn’t change with [T] (See Fig. 3D, inset and Section 6 in Materials and Methods) even though the mean lengths are very different for the two models for [T] above [*T*]_*c*_. This serves as a theoretical prediction for severing controlling length and can be used to discern this mechanism from other control mechanisms(Mohapatra et al., 2016).

## Discussion

Dynamics of protein filaments in cells (microtubules and actin filaments) are very different from what is observed in vitro (Zwetsloot et al., 2018). For example, actin networks in cells disassemble on the order of seconds whereas in vitro actin filaments disassemble over minutes (Miyoshi and Watanabe, 2013; Watanabe and Mitchison, 2002). These observations suggest that the dynamics of filament assembly are affected by proteins that associate with filaments in vivo and exert control over assembly and/or disassembly (Kuo and Howard, 2021; Sharp and Ross, 2012; Zanic et al., 2013). Inspired by recent experiments on microtubules (Vemu et al., 2018), we consider one such control system and demonstrate that it leads to dynamics that are distinct from those described for microtubules in isolation.

We model the severing of microtubules as a competition between the process of removal of monomers from the damage site created by a severing protein and the repair of the damage site by binding of free monomers. We use a one-dimensional random walk model to describe this competition and compute quantities that can be measured in experiments, such as the probability of severing and the time taken to successfully sever a microtubule as a function of the free tubulin concentration. We quantitatively compare our theory to a number of experimental observations(Vemu et al., 2018) pertaining to the effect of severing proteins on microtubules. Notably, we predict a distribution of severing times (Fig. 1D) skewed towards large severing times, which was observed in experimental results(Vemu et al., 2018). Additionally, using our model, we predict that the time to sever has a non-monotonic dependence on tubulin concentration and reaches a maximum at a critical tubulin concentration. Next, we studied the implication of this repair process on the length of a growing filament and found that it leads to a very sensitive dependence of steady state length as a function of free tubulin concentration.

### Signatures of severing as a length-control mechanism

Often in cells, several mechanisms can be at play to control the size of a sub-cellular structure, making it challenging to distinguish between them. Key results of this paper can serve as quantitative signatures of the severing mechanism.

A key feature of length-control mechanisms is that they lead to a peaked distribution of steady-state filament lengths. Several theoretical and experimental studies have demonstrated that severing can control the length of filaments (Kuo et al., 2019a; Mohapatra et al., 2016; Roland et al., 2008). Longer microtubules have more binding sites for severing proteins, and therefore have a greater probability of being severed than shorter ones. This leads to negative feedback on the length of a growing microtubule which eventually reaches a steady-state length once the disassembly rate balances the rate of assembly. The resulting distribution of steady-state lengths, in addition to being peaked, has a skewness (see Fig. 3D) which makes it distinct from other control mechanisms studied thus far (Mohapatra et al., 2016). This skewness, predicted in theoretical studies (Kuo et al., 2019a; Mohapatra et al., 2016; Tindemans and Mulder, 2010), was observed in recent experimental studies (Kuo et al., 2019a; Vemu et al., 2018). Additionally, we also find that the coefficient of variation (CV) of steady state microtubule lengths is independent of the concentration of tubulin and has a constant value equal to 0.523 (See Fig. 3D inset and Section 6 in Materials and Methods). This result relies on an assumption that the severing process is uniform along the filament, as was recently reported for spastin (Kuo et al., 2019a). Factors like filament aging, as has been documented in the context of actin filaments, arising due to the ATP hydrolysis (Roland et al., 2008), could invalidate this assumption.

### Experimental predictions

#### 1. Probability of severing as a function of tubulin concentration

We have shown that a one-dimensional random walk model quantitatively accounts for recent experiments on microtubule severing. Using this model, we find the probability of severing decreases as the concentration of tubulin increases (See Fig. 2B). This prediction of the model can be used to further test it. Many studies have used a “severing assay” to study the effect of severing proteins on microtubules (Belonogov et al., 2019; Vemu et al., 2018; Ziółkowska and Roll-Mecak, 2013). In these experiments, severing proteins are flown into a reaction chamber with pre-formed microtubules and quantities like the number of successful severing events per unit length of the microtubule (the severing rate) can thus be measured. Note that the maximal rate of severing is achieved when there is no free tubulin to repair the initial damage caused by a severing protein. Therefore, the ratio of the severing rate in the presence of free tubulin to the severing rate with no free tubulin is the probability of severing. At zero concentration of free tubulin, we expect that all severing initiation events lead to successful severing, but that the proportion would decrease as free tubulin can lead to repair, which is captured by equation 2, and Fig. 2B.

#### 2. Non-monotonic relationship of severing time and free tubulin concentration

Importantly, our model predicts a critical tubulin concentration [*T*]_*c*_ = *k*_*r*_/*k*_*T*_, where *k*_*r*_ is the first order rate of removal and *k*_*T*_ is the second order rate of addition of tubulin from the damage site. Above this critical concentration, the probability of severing drops rapidly, causing the severing time to peak at [*T*]_*c*_; see Fig. 2A. By using our estimated parameters, we predict that the mean severing time should increase for 0 ≤ [T] ≤ 5μM, a prediction which agrees with results from experiments described in a recent study(Vemu et al., 2018). However, at tubulin concentrations above those which were experimentally investigated in (Vemu et al., 2018), [*T*]>5 *μM*, we find that severing time decreases. This prediction can be used to test the proposed model using the set-up described in (Vemu et al., 2018).

#### 3. Ultra-sensitivity of microtubule length control

Severing of filaments, which are also growing by the addition of monomers, leads to length control. We observe that the tubulin repair process leads to an ultra-sensitivity in the lengths of microtubules as a function of tubulin concentration (Fig. 3C).

In Fig. 4, we show results of calculations of the change in steady-state length (*L*_*S*_) and the coefficient of variation as a function of two experimentally tunable knobs – tubulin and severing protein concentrations. We find a sharp increase in *L*_*S*_ with [*T*] above the critical concentration 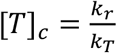 while the coefficient of variation remains the same across all concentrations of severing proteins. These results can be tested by creating an in vitro system (Díaz-Valencia et al., 2011; Kuo et al., 2019b; Vemu et al., 2018) in which growing microtubules are exposed to severing proteins. We also compute the probability distribution of steady-state lengths as a function of tubulin concentrations which can be used as a more stringent test of the model (see Fig. 3D).

**Figure 4:**
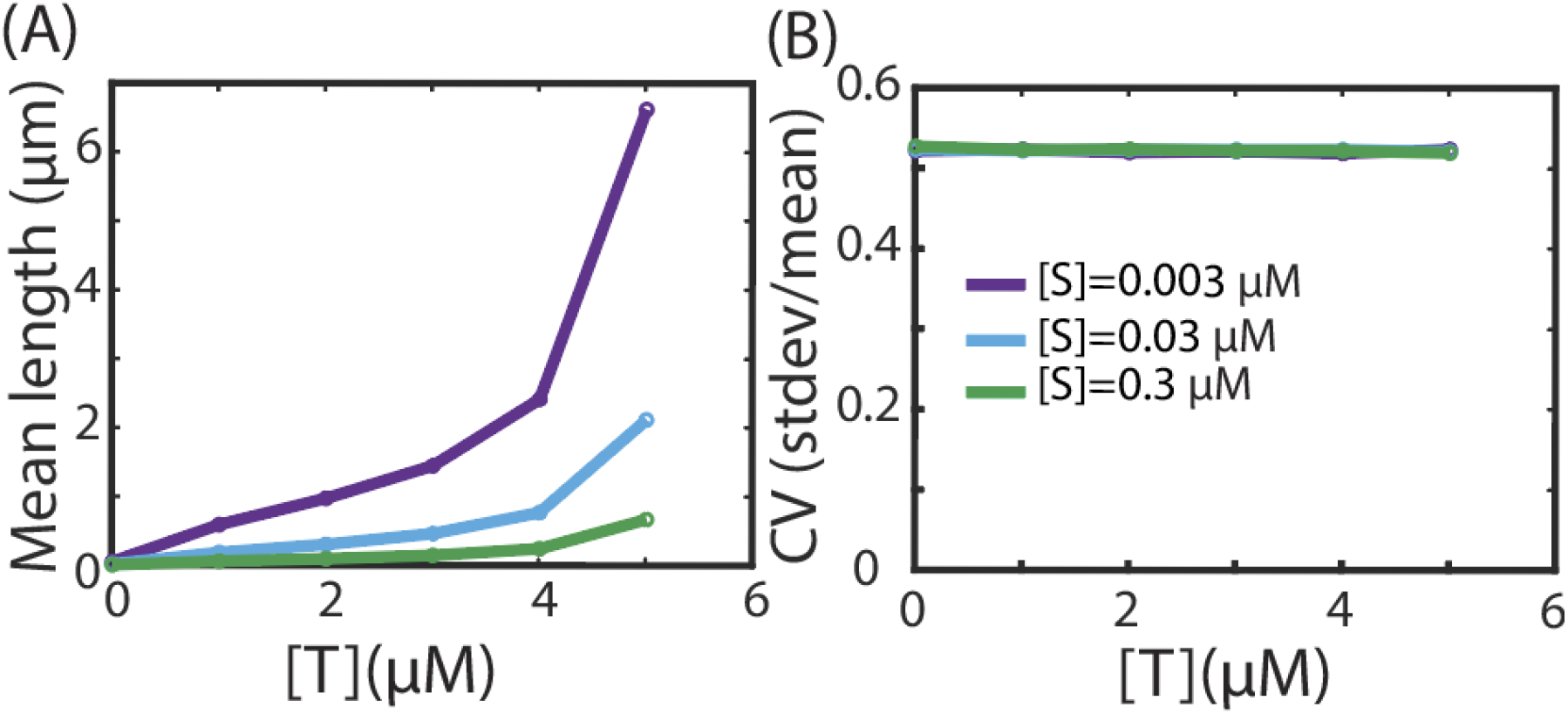
Testable predictions of the severing with repair model of length control. (A) Mean filament length and (B) coefficient of variation at steady state as function of Tubulin concentration at different severing protein concentrations. Dynamic range is larger at lower concentrations of severing proteins, and the coefficient of variation is independent of tubulin and severing protein concentrations. Parameters used in the simulation are listed in Table 1.

### Examining the assumptions of the model

In order to make the model of severing with repair analytically tractable, we made a few simplifying assumptions. Below we examine how robust our conclusions are with respect to these assumptions:

### Effect of finite severing time on microtubule dynamics

In our calculations of microtubule length-control (Figs. 3 and 4), we ignore the time it takes for the severing event to complete once the initial damage is created. This is equivalent to assuming that the time between subsequent damage creation events is much bigger than time to sever. This is a reasonable assumption in the limits of low and high tubulin concentrations, but not a valid one for intermediate tubulin concentrations (Fig A3, section A1 in Appendix).

Using the rates in Table 1, for small tubulin concentrations [*T*] ≈ 0.01*μM*, typical microtubules lengths are small, about 50 nm (using equation 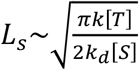 at small [*T*] approximation) and *P*_*sev*_[*T*] is close to 1, which means practically all damage events lead to severing. Time taken for a severing protein to bind to a microtubule and induce damage is 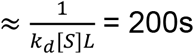 for a microtubule of length around 50 nm. In comparison, the severing time after initial damage 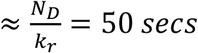. Thus, at low tubulin concentrations, severing time is shorter than the severing protein binding time, making it unlikely that multiple damage sites are created. At high tubulin concentrations[*T*]≳5 *μM*, the microtubules are ≳1 micron in length, and the time for a severing protein to bind is around 10 secs. At these tubulin concentrations, the probability of severing is small (≲0.025) and at least 40 binding events are required for one severing event. Waiting time for 40 binding events is about 400 seconds, which is greater than the severing time (200s or less) (Fig. A3 D, inset). Thus, our simplified model that ignores the possibility of multiple damage sites, in addition to being analytically tractable, provides a good approximation to study the effect of severing on microtubule length control in the limits of low and high tubulin concentrations.

At intermediate tubulin concentrations, however, time between severing events is comparable to severing time (Fig A3 D, inset), making it possible that several damage events are happening at once. This can lead to correlations between severing events which our calculations ignore. In order to study the implications of this scenario, we performed stochastic simulations wherein we tracked the locations of multiple damage sites on a microtubule lattice. Each damage site has a corresponding severing time, drawn from a distribution of severing times obtained from the stochastic simulation of the damage spreading simulations (described in Section 5(A) in Materials and Methods). We observe that results of large dynamic range in steady-state length from this simulation are similar to what was seen with the simple model (Figs. 3C, A4). Additionally, the noise as a function of free tubulin concentration from the full simulation was identical to the one obtained from the simple model at low and high tubulin concentration. However, at intermediate values of tubulin concentrations, we notice a peak in the CV vs tubulin concentration plot (Fig. A3 D), which was in contrast to the constant value seen with the simple model (Fig. 3D) but to be expected due to correlations in severing events.

### Dynamic instability

Microtubules are known to switch between phases of slow growth and rapid shrinking by a process known as dynamic instability. The rates of switching from shrinking to growth and vice versa are called rates of rescue and catastrophe, respectively. In our simulations leading to Fig. 3 and 4, we assumed that the microtubule are stabilized, and are not undergoing dynamic instability. In order to check the robustness of our results, we included dynamic instability in our simulations (see section A2 for simulation protocol). We found that dynamic instability did not qualitatively change the observation of large dynamic range of filament lengths. Coincidentally, a previous theoretical study analyzed the effect of the severing protein spastin on dynamic microtubules, and reported that their model predictions were similar to a model which ignored the dynamic instability (“no catastrophe” model) (Kuo et al., 2019a).

With stabilized microtubules, we had calculated that the coefficient of variation (CV) for the steady state length fluctuations for the models of severing without repair and severing with repair was constant with respect to tubulin concentration. In contrast, when dynamic instability was included, we observed a non-monotonic relationship in CV for both models (see Fig. A4 in section A2). In our simulations, microtubule filaments grow at a rate proportional to tubulin concentration. Since severing depends on microtubule length, which is typically small at low concentrations of tubulin, dynamic instability dominates over severing in both models. In this regime, the length distributions are exponential and resulting in CV being close to 1(Dogterom and Leibler, 1993; Kuo et al., 2019b). However, we observe different behaviors for both models (severing without repair and severing with repair) at higher tubulin concentration. In this regime, filaments grow longer and severing dominates over dynamic instability, as the number of subunits lost in a catastrophe event is small compared to the number of subunits lost due to severing. So, as expected, in the model of severing without repair, CV converges towards 0.523 at large concentration of tubulin. In severing with repair, we observe that CV shows a non-monotonic behavior with tubulin concentration, where it first decreases and then increases, and the dip in CV depends on the rescue rates chosen. Additionally, in-vitro studies have reported an increase in rescue rates in presence of severing proteins. Several hypotheses have been proposed to explain this, such as, the incorporation of new tubulin creating a “GTP-island”, which promotes rescue (Vemu et al., 2018) and the accumulation of severing proteins on the tips of shrinking microtubules, where they slow shrinkage and promote rescue (Kuo et al., 2019b). At this point, it is unknown whether severing proteins have the same effect in vivo, where they may interact with other microtubule associated proteins.

In summary, our calculations show that tubulin and severing proteins can cause qualitatively new dynamics of microtubules. Using simulations and theory, our study describes the role of a free pool of building blocks on the process of repair of damage created by severing proteins and its implications for the control of length of microtubules. Here we show the existence of a critical tubulin concentration above which severing becomes strongly dependent on [*T*], leading to a rare but fast severing. We find that microtubule length control via severing with tubulin-induced repair is ultra-sensitive to changes in the concentrations of tubulin leading to a dramatically increased dynamic range of filament lengths at steady state. This mechanism may have biological significance in processes like mitotic spindle size scaling during development where relatively small changes in microtubule growth rates have been correlated with large changes in mitotic spindle size(Rieckhoff et al., 2020). Our study shows how the antagonistic action of assembly and disassembly factors can produce novel dynamical properties of microtubules.

## Material and Methods

### 1. Calculation of probability of severing

Severing protein lands and creates a damage on a 1-dimensional grid of size *N*_*D*_. The fate of the damage is determined by a competition between the removal and addition of tubulin at the damage site, eventually leading to severing or repair as shown in Fig. A2 (A) in the Appendix. We assume that monomers are added to the damage site at a rate proportional to the concentration of free tubulin *k*_*T*_[*T*] (*s*^−1^) and are removed from the damage site at a rate *k*_*r*_ (*s*^−1^), with each step either increasing or decreasing the size of damage. If the size of the damage reaches a size *N*_*D*_, the filament is severed. On the other hand, if the size reaches zero, then the damage is repaired. Thus, the size of the damage *x* measured in monomers (i.e. tubulin dimers) is modeled as a one-dimensional random walk with absorbing boundaries at *x* = 0 and *x* = *N*_*D*_.

Probability of severing *P*_*sev*_(*x*) is the probability that 1-D random walker, starting at position *x*(0 ≤ *x* ≤ *N*_*D*_) reaches the *x* = *N*_*D*_ boundary before repair occurs, i.e., before *x* = 0 is reached. In this case, the probability of severing when the damage size is 0, i.e., *P*_*sev*_ (*x* = 0) = 0 and probability of severing when the damage size is *N*_*D*_ i.e., *P*_*sev*_(*x* = *N*_*D*_) = 1. These two conditions will serve as our two boundary conditions.

If 0 < *x* < *N*_*D*_, then, we can write the following recursion equation

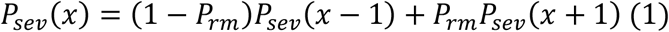

where *P*_*rm*_ = *k*_*r*_/(*k*_*r*_ + *k*_*T*_[T]) is the probability of damage spot increasing in size (monomers being removed from the site) and 1 − *P*_*rm*_ is the probability of damage spot decreasing in size (monomers being added at the site).

We use the method of characteristic equations to solve the recursion equation. Substituting, *P*_*sev*_ (*x* − 1) = 1, *P*_*sev*_ (*x*) = *s* and *P*_*sev*_ (*x* + 1) = *s*^2^, we get

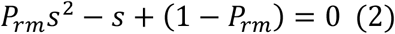

whose two solutions are 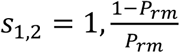. Solutions to (1) are *P*_*sev*_ (*x*) = *A*(*s*_1_)^*x*^ + *B*(*s*_2_)^*x*^. Substituting, we get 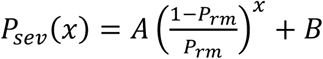. Using boundary conditions, *P*_*sev*_ (0) = 0, *P*_*sev*_ (*N*_*D*_) = 1, we get B = −A and 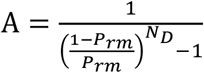. We define a dimensionless tubulin concentration 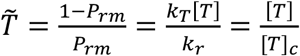 which is the ratio of the tubulin concentration and a critical concentration [*T*]_*c*_, which is the tubulin concentration where the rate of repair is equal to rate of severing, equal to *k*_*r*_/*k*_*T*_,. We find that 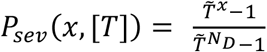, which is a function of Tubulin concentration. We note that when 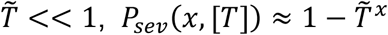 and when 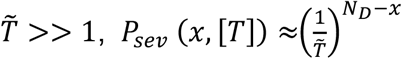. Also, 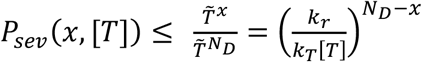, hence as the tubulin concentration increases, *P*_*sev*_(*x*) decreases (Fig. 2 and A2).

If we assume an initial damage spot of a size *x* = 1 as we do in the manuscript, then 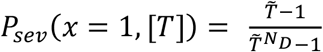 which is approximately equal to 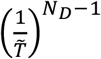 when 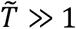 and 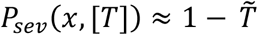 when 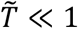. In other words, above the critical tubulin concentration, the probability of severing becomes a very sensitive function of the tubulin concentration.

### 2. Calculation for time to either sever or repair

Let 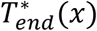 be the average time (in steps) it takes to for the filament to either get severed or repaired, starting with a damage of size *x*. On the first step, the damage spot can either get replenished with monomers with a probability 1 − *P*_*rm*_ or increase in size by having monomers removed with a probability *P*_*rm*_ in time Δ*t* i.e.

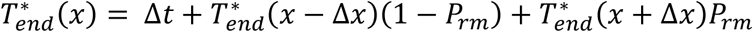

There are two possible boundary conditions, 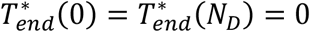. Using Δ*x* = Δ*t* = 1, we get a recursion relation similar to the one before, i.e. 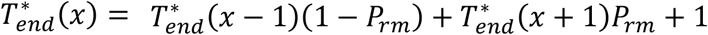, which is an inhomogeneous recursion relation.

We first find the solution to the homogenous equation:

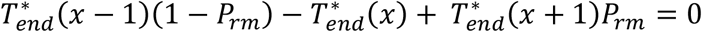

This is similar to the one we solved in Section 1 and following the same method, we can show that the solution is 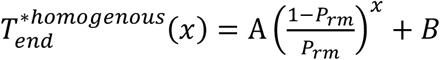 and the solution to inhomogeneous equation we have to guess (since the inhomogeneous is a constant) 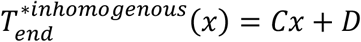

Substituting it in the equation for 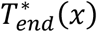 we get 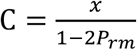 and hence the complete solution is 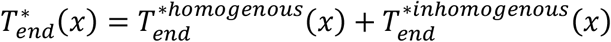 which is 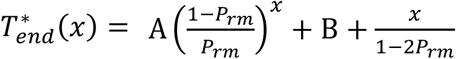

Using the boundary conditions, and assuming *P*_*rm*_ ≠ 1/2, we find 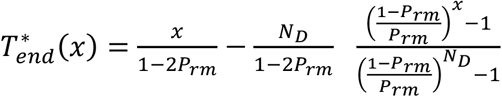. When *P*_*rm*_ = 1/2, 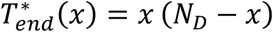.

Note that 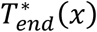 is the time (in steps) taken to reach either end (filament gets repaired or severed). We get time taken by a damage spot to be repaired or severed, i.e. *T*_*end*_(*x*) by multiplying time in steps by 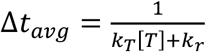,

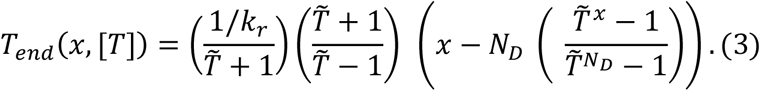

where 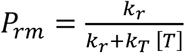 and 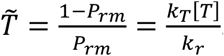. Equation 3 for (*x* = 1) is plotted in Fig A1 (C).

### 3. Calculation of severing time

In this section, we derive an expression for time it takes to sever (equation 1 in main text) a filament. In the one-dimensional random walk model for damage spreading, there are two ways to reach the end – either get severed or get repaired. Hence, the time to reach either end is given by the following equation,

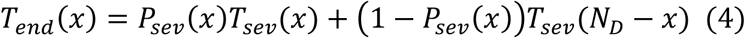

where *T*_*end*_(*x*) is the time to reach either end if the initial size of the damage is *x, T*_*sev*_(*x*) is the time to sever and *P*_*sev*_(*x*) is the probability to sever when starting at *x*. Using the expression for *T*_*end*_(*x*) which we computed in Section 2 (Materials and Methods), we can extract *T*_*sev*_ (*x*).

Replacing *x* with *N*_*D*_ − *x* in equation 4 we get, *T*_*end*_(*N*_*D*_ − *x*) = *P*_*sev*_(*N*_*D*_ −*x*)*T*_*sev*_ (*N*_*D*_ − *x*) + 1 − *P*_*sev*_ (*N*_*D*_ − *x*) *T*_*sev*_(*x*). Using 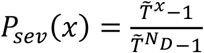 and 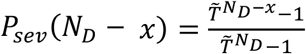, we get 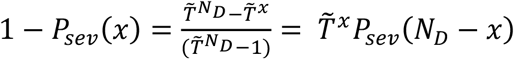 and 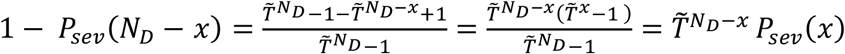.

From section 2 in Material and methods, 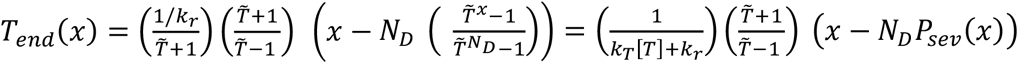.

Substituting the expression for 1 − *P*_*sev*_(*x*) and 1 − *P*_*sev*_(*N*_*D*_ − *x*) in the equations for *T*_*end*_ (*x*) and *T*_*end*_ (*N*_*D*_ − *x*), we get

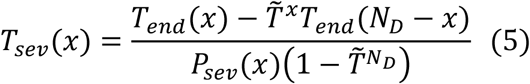

After substituting expression for *T*_*end*_ (*x*) and *T*_*end*_ (*N*_*D*_ − *x*) in (3), we get

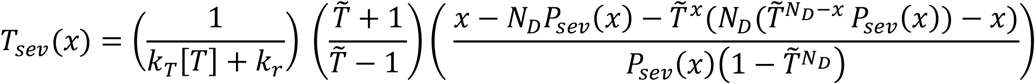

Finally, after substituting the expression for *P*_*sev*_(*x*) and moving terms around, we get

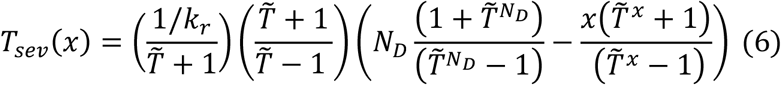

For an initial damage size 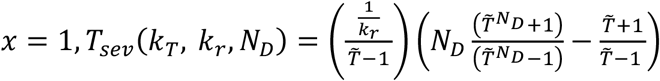, which is the equation we used to plot the mean severing time in Fig. 1D and A2(C) inset (see Appendix).

*T*_*sev*_ (*k*_*T*_, *k*_*r*_, *N*_*D*_) has two limits: (1) When 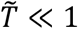, and 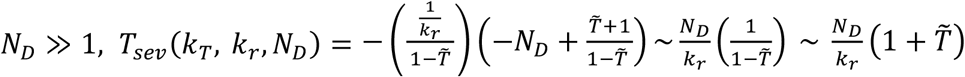 the severing time is linear in tubulin concentration. (2) When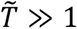, and 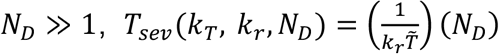 the severing time decreases as tubulin concentration increases. These limiting behaviors imply that the severing time has a maximum as a function of tubulin concentration, which is obtained at a concentration that is of the order critical tubulin concentration 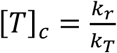.

For the diffusive limit, when the rate of damage spreading and repair are balanced i.e. when 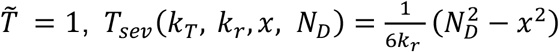.

### 4. Estimation of parameters

In this section, we compare our theoretical predictions with experimental results from ref. (Vemu et al., 2018), and extract values for the parameters of our model. In particular we focus on Fig. 3D and 3H in ref (Vemu et al., 2018), where the authors plot probability to sever and time taken to sever for different concentrations of tubulin.

Our model for severing with repair has four parameters - *k*_*T*_, *k*_*r*_, *x* and *N*_*D*_. In equation 6, we find an expression for the mean severing time as a function of the tubulin concentration. We assume an initial damage of size *x* = 1, and constrain the space of possible parameter values using experimental data in ref (Vemu et al., 2018). In equation 2, we predict that the probability of severing decreases as *N*_*D*_ increases (Fig. A1). We use the observation of severing events at [*T*] = 5 *μM* in ref (Vemu et al., 2018) to constrain *N*_*D*_ (upper bound = 40) by assuming that probability of severing is at least 10^−3^. We obtain a rough lower bound for *N*_*D*_ ≈ 20 from the TEM images in Fig. 1 in ref. (Vemu et al., 2018) showing patches of tubulin removed from microtubules which are still intact. Next, we use the method of least squares on this constrained parameter range to find sets of parameters that fit the published data Fig. 3H in ref (Vemu et al., 2018) which reports the mean and the standard deviation of severing times as a function of tubulin concentration (Fig. A2). Our estimated values are: *k*_*T*_ = 0.11 *μM*^−1^*s*^−1^, *k*_*r*_ = 0.54 *s*^−1^ and *N*_*D*_ = 25. Using these estimated values, we plot the severing time and the probability of severing in Fig. 2A and 2B respectively.

#### Sensitivity analysis of parameters

As seen in Fig. A1, probability of severing increases with a larger initial damage size *x* but severing time is unaffected. Probability of severing is unaffected by *N*_*D*_ but time to sever increases with a larger *N*_*D*_. As is captured by the analytical formula in equation 2 in main text, the critical concentration of tubulin [*T*_*c*_] where the time of severing starts decreasing, is unaffected by *x* and *N*_*D*_.

#### Comparison of experiment and theory via bootstrapping

In Figure 1 in the main text, we show a comparison of experimental data from (Vemu et al., 2018) with a theoretical curve for mean severing time and simulation curve for standard deviation of severing time using our fitted parameters. Since ref(Vemu et al., 2018) reported mean severing time and standard deviation of severing time, we used a bootstrapping scheme using data resampling to compute the error on the mean (error bars in Fig 1C), and error on the standard deviation (error bars in Fig 1D(inset)). The error bars on the mean severing time and standard deviation of severing times were calculated from 750 bootstrapping samples.

### 5. Stochastic simulation schemes

#### (A) Model for severing with repair

A severing event begins when a severing protein lands on a filament, binds to the side, and creates a damage of size *x*. The fate (eventual severing or repair) of the damage created by a severing protein is determined by a competition between the removal and addition of tubulin at the damage site. We use the Gillespie algorithm to simulate this process - monomers are added to the damage site at a rate proportional to the concentration of free tubulin *k*_*T*_[*T*] (*s*^−1^) and are removed from the damage site at a rate *k*_*r*_ (s^-1^), with each step either increasing or decreasing the size of damage. If the size of the damage reaches a size *N*_*D*_, the filament is severed. On the other hand, if the size reaches zero, then the damage is repaired. By performing this simulation multiple times (typically about 13,000 severing events), we obtain, as a function of tubulin concentration, the probability of severing (Fig. 2), the mean severing time (Fig. 1C) and the probability distribution of severing times (Fig. 1D).

#### (B) Length trajectory with repair-severing

Simulation of filament growth is performed using a Gillespie algorithm. At each step of the simulation, either the filament gains one subunit, with probability proportional to the concentration of tubulin (*k* [*T*]), or it gains a new severing site when a severing protein lands on the filament at position between 1 and *l* − 1, with probability *k*_*r*_, where *l* is the length of the filament. In the absence of competition between severing and repair, a severing event occurs immediately upon creating. However, if competition is considered, each time a severing site is created, a severing event occurs at this site with a probability *P*_*sev*_(*x*) in equation 1.

#### (C) Filament growth with finite severing time

In order to incorporate a finite severing time into our simulation, we first run our damage spreading simulation (A) to create a distribution of severing times. Then, we adjust the filament growth simulation in (B) by tracking the location of each damage site (where severing protein binds) on our microtubule. The severing occurs with a probability given by equation (2). For every severing event, we assign a severing time by sampling it from our distribution of severing times. Once this time has surpassed in the simulation, the microtubule is severed at this damage site. We keep track of one part of the severed microtubule. The other part, and any damage sites on that part, are removed.

### 6. Calculation of probability distribution of filament lengths, due to severing with and without repair

In ref(Mohapatra et al., 2016), we considered the case of a uniform rate of severing along a filament; in other words, severing takes place anywhere along the filament with equal probability. A filament consisting of *l* subunits can be broken into two smaller filaments at any of the (*l* − 1) positions with an equal rate *s*, for any choice of severing location; the total rate of severing at any location is then, *s* (*l* − 1). In addition to the severing rate, there is *r*, the rate at which subunits are added to a filament. Using these two rates we find an expression for the probability distribution of lengths at steady state (reproduced from ref(Mohapatra et al., 2016)), 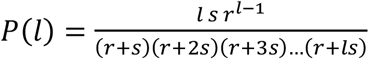, which can be further simplified to

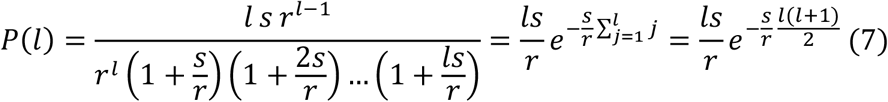

Here we use the approximation that 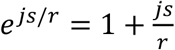 which is valid for *js* ≪ *r*. For 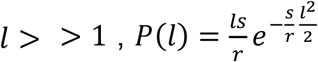 and we use this expression to plot the distribution of lengths without repair with rates *r* = *k*[*T*] and *ss* = *k*_*d*_[S], and with repair with the rates *r* = *k*[*T*] and *s* = *k*_*d*_[S]*P*_*sev*_([*T*]) in Fig. 3D.

Additionally, we can use this approximation to compute the mean < *l* > and variance of the filament length distribution, < *l*^2^ > − < *l* >. In general, moments are defined as, 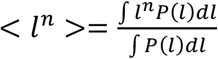. For the mean, *n* = 1, and hence 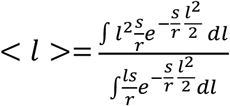.

Using the properties of gaussian integrals, 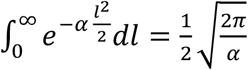, we can show 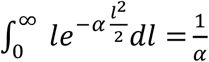, and 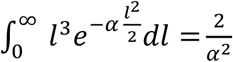. Substituting these results, we get 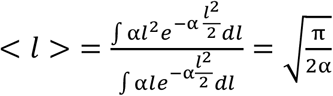 and 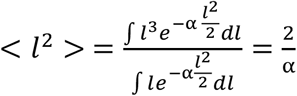.

The variance, < *l*^2^ > − < *l* >, is 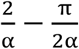 and standard deviation, 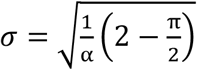.

We define “Noise” or the coefficient of variation as 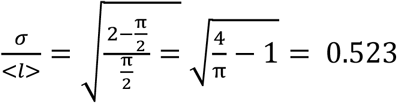, which is a prediction for the uniform severing mechanism. We use this relationship to plot Fig. 3D, inset. Additionally, substituting 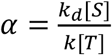 in to the expression for the mean and variance of filament lengths at steady state, we get, 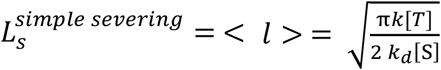, and substituting 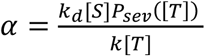, we find 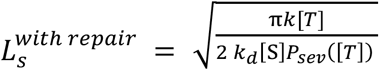, which is used in Fig. 3C.

Note that 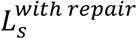 has two regimes: When 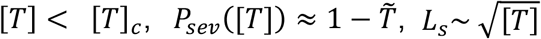. When 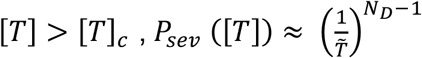 and, 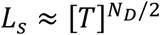.

## Appendix

**Fig A1.**
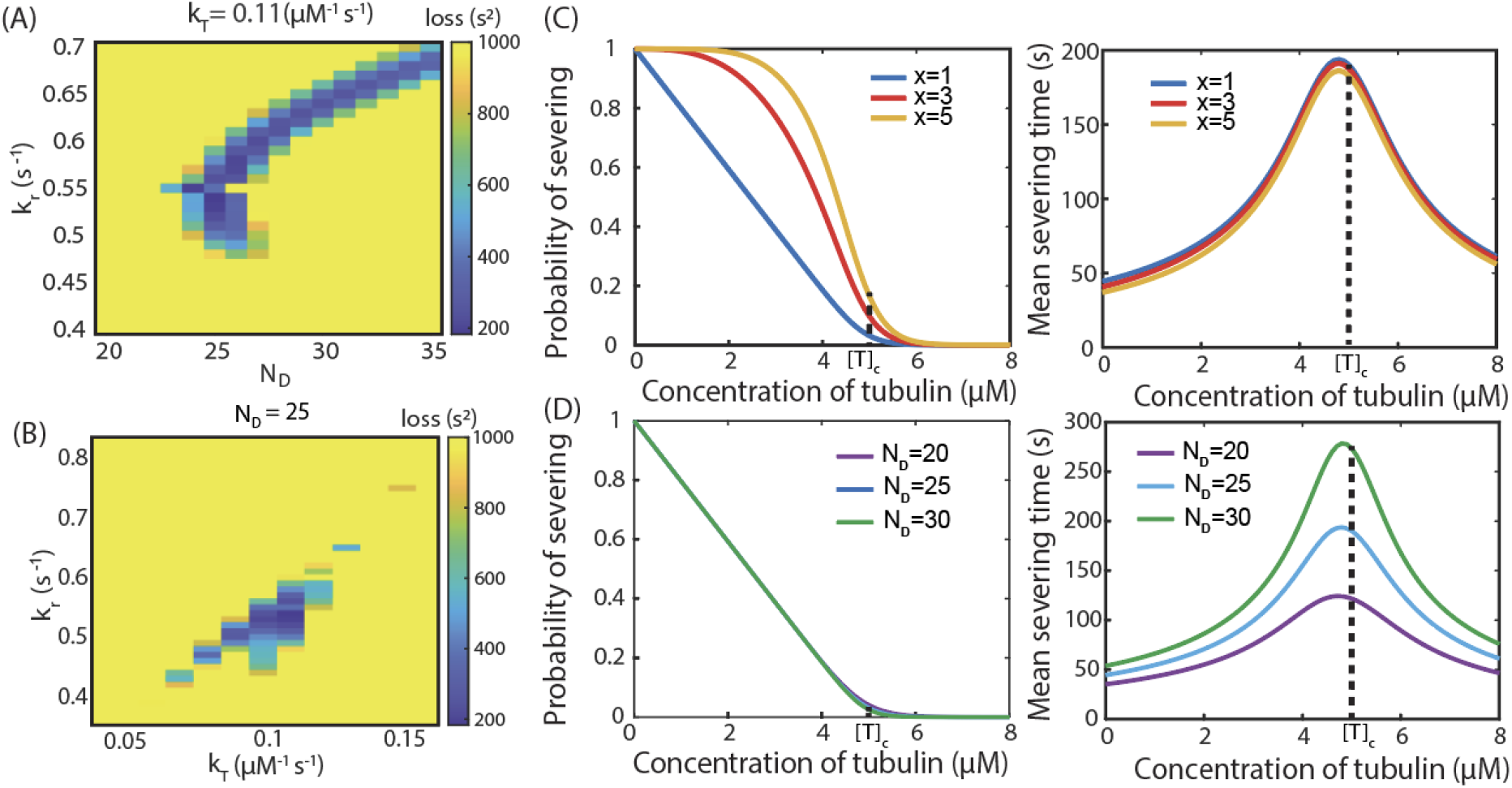
Estimation of parameters. (A) and (B) Density plots for the estimation of ***k***_***r***_, ***k***_***T***_ and ***N***_***D***_. Loss function calculates the sum of squares of differences between mean severing time from equation (6) with indicated parameters and experimental data for [T]=0,2,5 μM. Probability of severing and Mean severing time vs concentration of Tubulin as a function of (C) ***x*** and (D) ***N***_***D***_.

**Fig A2.**
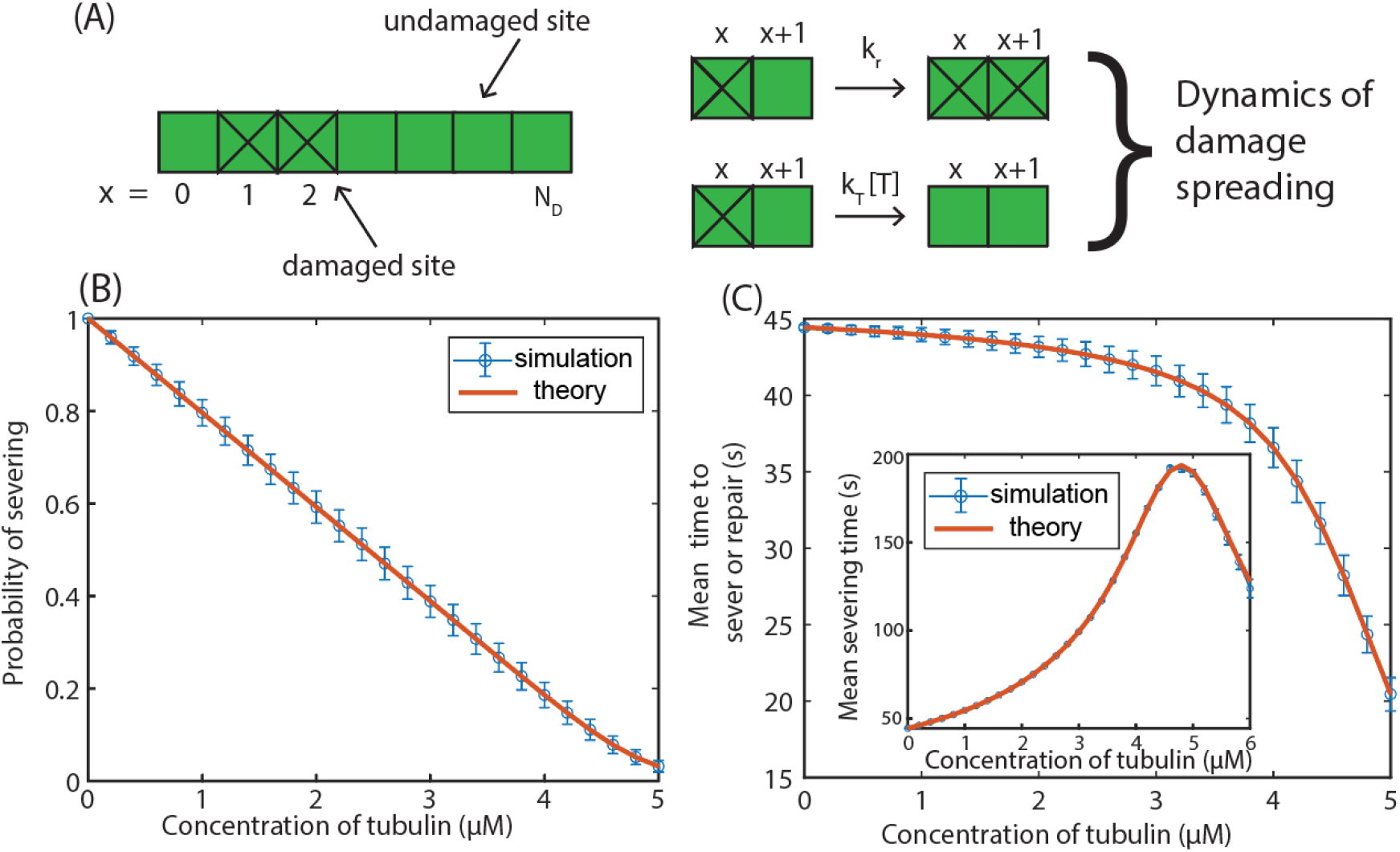
Comparison of theory and simulation for severing with repair. (A) Schematic of damage spreading in a damage site, corresponding to Figure 1B in the main text. Damage spreads along the microtubule lattice, i.e., tubulin dimers are removed, at a rate ***k***_*r*_, and damage is repaired, i.e. tubulin dimers are incorporated back into the microtubule lattice a rate ***k***_***T***_[***TT***]. (B) Probability of severing and (C) time to repair or sever and mean severing time (inset) as a function of tubulin concentration. All plots were computed using 3000 damage spreading simulations, and the SEM error (in blue) was calculated by repeating these 2000 times. Parameters used are listed in Table 1.

### A1: Finite severing time

If the severing protein binding time is comparable to severing time, it is possible that, after one severing event, there are other damage sites on the filament which will lead to a severing event shortly thereafter. This will produce correlations in subsequent severing events which we ignore when deriving our analytic formulas for *P*_*sev*_ and *T*_*sev*_. In order to account for this situation, we performed a simulation where we allowed multiple damage sites on a microtubule lattice at one time, see Section 5(C) in Materials and Methods. We observed that our results of large dynamic range in steady-state length from the full simulation are similar to what was seen with the simple model where we assumed a negligible severing time (Fig. A3). The coefficient of variation (CV) of filament lengths at steady state calculated in the full simulation was identical to the one obtained from the simple model at low and high tubulin concentration. However, at intermediate values of tubulin concentrations, we observed a peak in the CV as seen in Fig. A3(C), which was in contrast to the constant value seen with the simple model (Fig. 3D). We observed that the peak in CV depends on the concentration of severing protein as shown in Fig. A3(D). As shown in inset, the variation from the typical constant value occurs when the time between binding events which can lead to severing (computed by 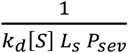 where *L*_*s*_ is the length of microtubule) is less than the time it takes to sever.

**Fig A3.**
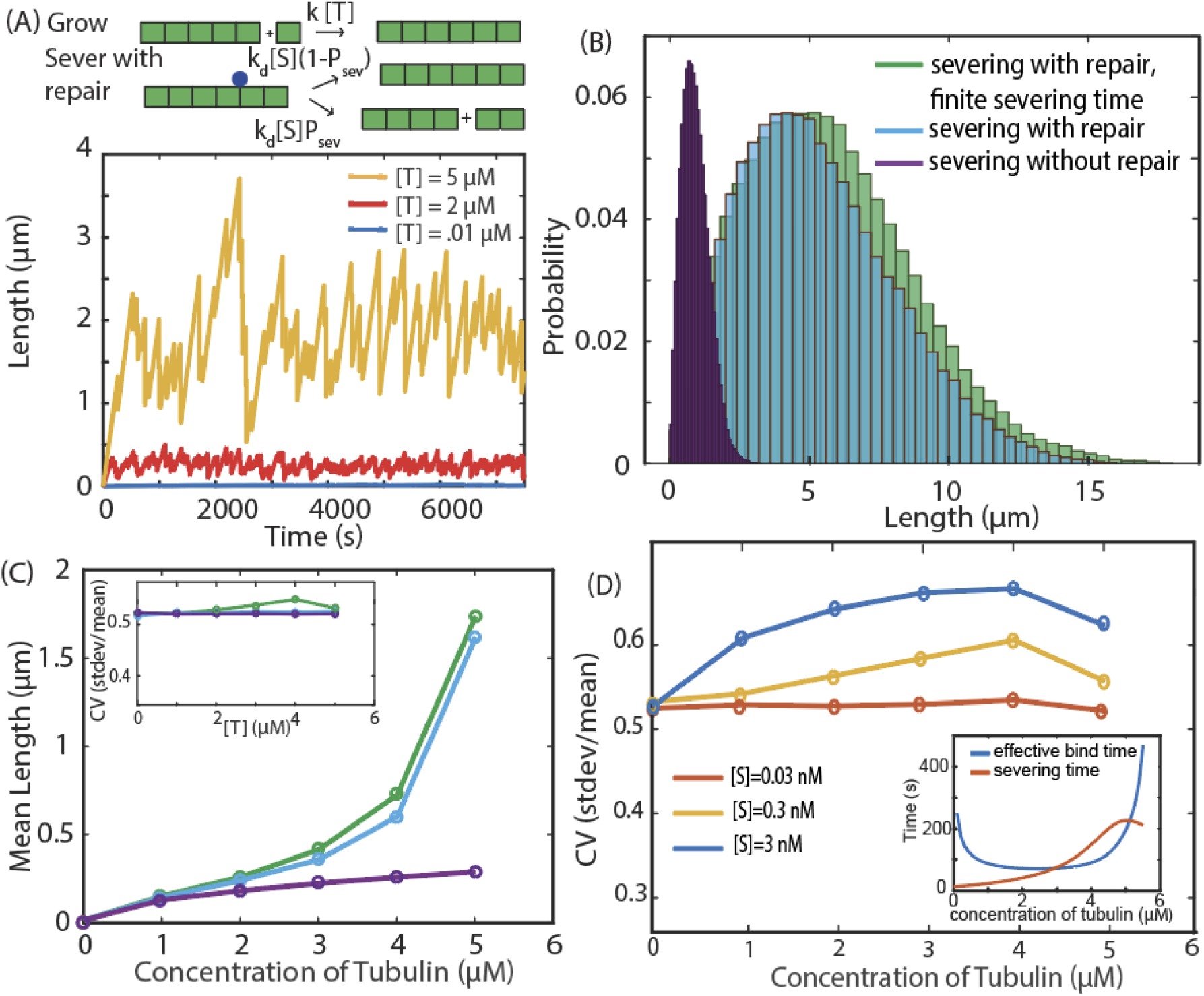
Effect of finite severing time on filament dynamics. (A) Length trajectory of a microtubule (average of three for clarity) which takes into account the finite time it takes to sever a filament, at different concentrations of tubulin. (B) Probability distribution of steady state lengths, computed using 10^6^ trajectories. (C) Mean steady state length and coefficient of variation (CV) (inset) with and without repair (with and without finite severing time), computed using 10^6^ trajectories. (D) Adding a finite time of severing does not change the results qualitatively for mean length, but there is a peak in the CV that depends on the severing protein concentration; the CV was computed using 3 × 10^6^ trajectories. (Inset) The CV differs from its constant value at tubulin concentrations at which the time between productive events that lead to severing (“effective bind time”) is less than the severing time. Effective bind time is given by 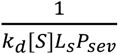. Parameters used in all simulations are listed in Table 1.

### A2: Dynamic instability

The simulation that includes dynamic instability follows the same algorithm as that detailed in Section 5(B) in Materials and Methods, with inclusion of an additional boolean parameter for whether the microtubule is in a state of rescue or catastrophe. In the rescue state, the microtubule grows (gain subunits) at a rate proportional to the tubulin concentration, *c*_*T*_[*T*]. During catastrophe, the microtubule loses subunits at a rate *C*_*s*_. The microtubule can undergo severing in either state. At each timestep, the microtubule can a) grow or shrink depending on whether it is in catastrophe or rescue, b) have a severing protein land and either sever the filament or not depending on the outcome of the competition between severing and repair, or c) switch from catastrophe to rescue or rescue to catastrophe. We use the parameters listed in Table 1 and A1 for our stochastic simulations and are in the regime where the growth/shrinking rates are faster than the rescue/catastrophe rates. As before, we find that the dynamic range of the mean length of microtubules is greater when the repair process is included (Fig. A4). However, coefficient of variation shows a non-monotonic relationship with tubulin concentration.

### A3: Conversion between monomer number and filament length

Our simulations are based in subunits. We use the following conversion to convert from subunits to length: (4/13) (length in # of subunits) = (length in nm). Microtubules are hollow cylinders and made up of 13 protofilaments built up from tubulin dimers, each 4 nm in length. A cross section of a microtubule is a rind of 13 dimers that is 4 nm wide.

**Fig A4.**
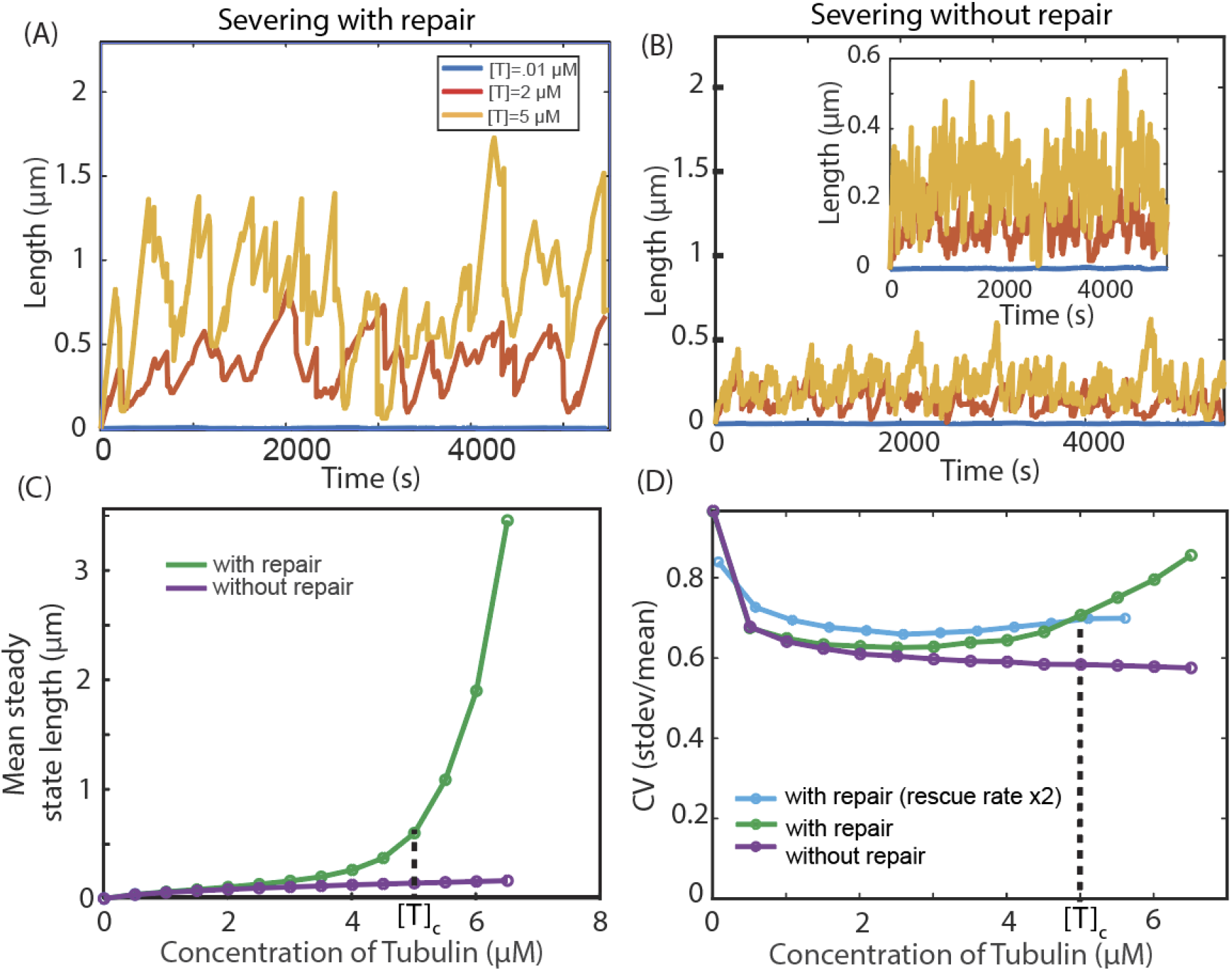
Effect of Dynamic Instability. (A) Length trajectory (average of 3) with repair and (B) without repair. (C) Mean steady state length, computed using 10^6^ trajectories and (D) coefficient of variation without and with repair with two rescue rates, computed using 10^6^ trajectories. Parameters used are listed in Table A1.

**Table A1.** Parameters use for studying Length control in presence of repair with dynamic instability.

| Parameters | Value used in paper | References |
| --- | --- | --- |
| Catastrophe rate | .004 s<sup>-1</sup> | (Margolin et al., 2012) |
| Rescue rate | .02 s<sup>-1</sup> | (Margolin et al., 2012) |
| Shrinkage rate in catastrophe | 152 dimers/s | (Margolin et al., 2012) |

## Acknowledgments

We wish to thank Stephanie C. Weber and Shane G. McInally for careful reading of the manuscript, and members of Mohapatra group and Kondev group for stimulating discussions. This work was supported by the National Science Foundation through grants DMR-1610737, MRSEC DMR 2011846 (J.K., C.S.), the Simons and the HHMI foundations (J.K., L.M. and C.S) and Rochester Institute of Technology start-up funds (L.M.). We acknowledge computational support from the Brandeis HPCC which is partially supported by the NSF through DMR-MRSEC 2011846 and OAC-1920147.

## Competing interests

Authors declare no competing interests.

## Notes

### Competing Interest Statement

The authors have declared no competing interest.

### Summary of Updates

In the revised version, the title was updated, along with some text in the Introduction.

